# The human small intestine contains two subsets of regulatory Foxp3+ CD4+ T cells with very different life span and functional properties

**DOI:** 10.1101/2020.02.12.941369

**Authors:** Sudhir Kumar Chauhan, Raquel Bartolomé Casado, Ole J.B. Landsverk, Hanna Johannessen, Danh Phung, Frank Sætre, Jørgen Jahnsen, Rune Horneland, Sheraz Yaqub, Einar Martin Aandahl, Knut E.A. Lundin, Espen S. Bækkevold, Frode L. Jahnsen

**Affiliations:** Department of Pathology, Oslo University Hospital-Rikshospitalet, Oslo, Norway; Department of Cancer Immunology, Institute for Cancer Research, Oslo University Hospital, Oslo, Norway; Institute of Clinical Medicine, University of Oslo, Oslo, Norway; Department of Gastrointestinal and Pediatric Surgery, Oslo University Hospital-Rikshospitalet, Oslo, Norway; Department of Gastroenterology, Akershus University Hospital, Lørenskog, Norway; Department of Transplantation Medicine, Section for Transplant Surgery, Oslo University Hospital-Rikshospitalet, Oslo, Norway; Department of Gastrointestinal Surgery, Oslo University Hospital - Rikshospitalet, Oslo, Norway; Department of Gastroenterology, Oslo University Hospital - Rikshospitalet, Oslo, Norway

**Keywords:** regulatory T cells (Tregs), CD4 T cells, human small intestine, Foxp3, transplantation, Helios, celiac disease

## Abstract

**Background:** Regulatory CD4 T cells (Tregs) in the mice gut are mainly specific for intestinal antigens and play an important role in the suppression of immune responses against harmless dietary antigens and members of the microbiota. However, information about the phenotype and function of Tregs in the human gut is limited.

**Objective:** Here, we performed a detailed characterization of Foxp3+ CD4 Tregs in the human small intestine.

**Methods:** Tregs and conventional CD4 T cells derived from normal intestine as well as from transplanted duodenum and celiac disease lesions were subjected to extensive immunophenotyping and their suppressive activity and ability to produce cytokines were assessed.

**Results:** Small intestinal Foxp3+ CD4 T cells were CD45RA- CD127- CTLA4+ and suppressed proliferation of autologous T cells. Approximately 60% of the Tregs expressed the transcription factor Helios. When stimulated, Helios- Tregs produced IL-17, IFN-γ and IL-10, whereas Helios+ Tregs produced very low levels of these cytokines. By sampling mucosal tissue from transplanted human duodenum we demonstrated that donor Helios+ Tregs have a short life span whereas Helios- Tregs persisted for at least 1 year post-transplantation. In normal small intestine, Foxp3+ Tregs constituted only 2% of all CD4 T cells, while in active celiac disease both subsets expanded 5-10-fold.

**Conclusion:** The small intestine contains two subsets of Tregs with different functional capacities and very different life span. Both subsets are scarce in the normal situation but increase dramatically in a chronic inflammatory setting.

## INTRODUCTION

The intestine is a challenging environment for the local immune system, which has to respond effectively to eliminate infectious pathogens while at the same time avoiding detrimental inflammatory responses to ubiquitous food antigens and the normal gut microbiome - a function termed oral tolerance. The mechanisms underlying oral tolerance are not clear, but it is thought that Foxp3+ CD4 T regulatory cells (Tregs) residing in the intestinal mucosa play a central role in this process ^1, 2^.

Our current knowledge about gut Tregs is mainly derived from mouse models. Tregs are found in most organs of mice, but those residing in the intestinal mucosa appear to have gut-specific phenotypes and functions ^2^. Whereas most tissue Tregs are directed towards self-antigens ^3^, intestinal Tregs express a T cell receptor (TCR) repertoire specific for food and microbiota antigens ^2^. The majority of Tregs in the mouse colon are microbial-specific ^4–6^, whereas most Tregs in the mouse small intestine (SI) respond to food antigens ^7^. Tregs in the intestine therefore appear well suited to regulate harmful immune reactions to luminal antigens ^1, 2, 7^.

Tregs specific for exogenous antigens originate from naïve CD4 T cells that are induced in the periphery (pTregs). Retinoic acid produced by intestinal dendritic cells induces both a Treg phenotype and upregulates gut homing receptors ^1, 8^. In contrast, Tregs specific for self-antigens are thymus-derived (tTregs) ^3^. It has been suggested that the expression of the transcription factor Helios can distinguish tTregs from pTregs, the former being Helios+ ^9^. However, in an elegant study analyzing the occurrence of Tregs specific for mucosal antigens in humans it was found that expanded memory Foxp3+ Tregs specific for grass, birch, mite and Aspergillus fumigatus all expressed high levels of Helios ^8^. Therefore, at least in humans, the expression of Helios appears not to distinguish tTregs from pTregs ^8, 10^.

The importance of Tregs in humans is documented by the fact that mutations of the Foxp3 gene are causative of the IPEX syndrome ^11^. This syndrome is characterized by a detrimental inflammatory state in many organs, including the intestine. The inhibitory receptor CTLA-4 plays an important role for the suppressive effect of Tregs, and colitis is the most prevalent adverse effect when anti-CTLA-4 monotherapy is given to treat cancer patients ^12, 13^. Together, these findings strongly suggest that Tregs are important for homeostasis in the gut mucosa. However, studies directly examining Tregs in the human gut are scarce, in particular in the SI.

Here, we performed a detailed characterization of Tregs in the human SI showing that these Tregs consist of two subsets – distinguished by their expression of Helios. The two subsets had distinct phenotypic and functional properties and displayed very different lifespan. Both subsets were scarce in SI compared with their colonic counterparts but increased dramatically in chronically inflamed tissue.

## MATERIALS AND METHODS

### Subjects and biological material

Duodenum- proximal jejunum resections were obtained from non-pathological SI during surgery for cancer in the pancreas or distal bile duct with a Whipple procedure (pancreatoduodenectomy) (n = 28, mean age 67 yr, range 50-84, 10 female) or from donor and recipient duodenum during pancreas transplantation of type I diabetes mellitus patients (donors: n = 13, mean age 33 yr, range 5-54, 5 female; recipients: n = 12, mean age 42 yr, range 25-53, 7 female ^14^). Endoscopic biopsies were obtained from donor and patient duodenum at 3, 6, and 52 weeks after transplantation. Colonic biopsies were obtained by colonoscopy of individuals with unexplained stomach pain with normal histology (n = 3, mean age 67 yr, range 63-72, all male) or non-tumor tissue from resections of colorectal cancer (n = 2, mean age 77 yr, range 71-83, 1 female). All samples were evaluated by an experienced pathologist and only material with normal histology was included. Duodenal biopsies were also obtained from newly diagnosed untreated celiac patients (n = 14, mean age 40 yr, range 19-74, 10 female). The study was approved by the Regional Committee for Medical Research Ethics in Southeast Norway and the Privacy Ombudsman for Research at Oslo University Hospital–Rikshospitalet and complies with the Declaration of Helsinki. All participants gave their written informed consent. Intestinal resections were opened longitudinally and rinsed thoroughly in PBS, and mucosal folds were dissected off the submucosa. For microscopy, small mucosal pieces were fixed in 4% formalin and embedded in paraffin according to standard protocols. To obtain single-cell suspensions, epithelial cells were removed by washing in PBS containing 2 mM EDTA three times for 20 min at 37°C, and the lamina propria was minced and digested in RPMI medium containing 2.5 mg/ml Liberase TL and 20 U/ml DNase I (both from Roche) at 37°C for 1 h. Digested tissue was passed through 100-μm cell strainers (Falcon) and washed three times in PBS. PBMC were isolated by lymphoprep gradient centrifugation of blood from patients or healthy buffycoats from the Oslo University Hospital Blood Center. Identically treated PBMCs (from healthy buffycoats, and patients) served as controls for the collagenase-sensitivity of epitopes.

### Flow cytometric analysis

Cells were stained with Fixable Viability Dye eFluor 780 (1 μl/10^6^ cells, eBioscience, San Diego, CA) for 30 min on ice, followed by surface staining with antibodies to CD3 (clone OKT3), CD4 (clone OKT4), CD8 (clone SK1) and CD127 (clone M21) for 15 min on ice. To detect intracellular cytokines, cells were treated with FOXP3/transcription factor staining buffer set according to the manufacturers protocol (eBioscience) and stained with antibodies to FOXP3 (clone 236A/E7), Helios (clone 22F6), and IL-17 (clone 64DEC) from eBioscience, or CTLA-4 (clone L3DTO), IFN-g (clone 4S.B3), and IL-10 (clone JES3-9D7) from Biolegend. All antibodies against intracellular/nuclear markers were incubated for 30 min on ice. For in-depth multicolor immunophenotyping of biopsies from celiac disease patients, single-cell suspensions were stained in bulk with a cocktail of surface backbone markers (CD45, CD3, CD4, CD8, CD69, CD103, CD69 and CD127) and followed by intracellular staining of Foxp3 and Helios using the FOXP3/transcription factor staining buffer. Then, cells were washed and aliquoted in a 96-well plate for incubation with a panel of different PE-conjugated antibodies. Information regarding all the antibodies used in the study can be found in the **supplementary Table 1**. For comparison, the same staining was applied to digested LP cells isolated from control samples (duodenal resections), in culture media or stimulated for 48h with α-CD3/CD28 coated beads (Dynabeads Human T-Activator, Thermo Fisher scientific) in a 1:1 ratio (beads to cells), using the manufactures protocol. Flow cytometry was performed on a BD LSRFortessa (BD Biosciences), and analyzed using FlowJo 10.3 and 10.6 software (Tree Star, Eugene, OR). Gates for surface markers were set based on fluorescence-minus-one (FMO) controls and irrelevant isotype-matched antibodies, and cytokines gates was based on untreated cells.

### Cytokine analysis

To assess the cytokine production by mucosal T cells, dispersed cells were cultured for 4 h in RPMI/10% fetal calf serum (FCS) with 1.5 ng/ml phorbol 12-myristate 13-acetate (PMA) and 1 μg/ml ionomycin (all from Sigma-Aldrich, St Louis, Mo) with Golgi-Stop (BD Bioscience, San Jose, CA) added after 1 h of stimulation to allow intracellular accumulation of cytokines.

### Treg suppression assay

High expression of CD25 is a hallmark of Tregs and we used CD25 to sort SI Tregs. However, surface CD25 was completely cleaved after enzymatic treatment during the sample preparation. We therefore first cultured tissue-derived single cells over-night and sorted Tregs after they had regained CD25 expression. Autologous CD25^−^CD127^+^CD45RA^+^ naïve CD4+ T cells were used as responder T cells (Tresp). Sorted Tregs were pre-activated for 48h using α-CD3/CD28 coated beads (Dynabeads Human T-Activator, Thermo Fisher scientific) in a 1:1 ratio (beads to cells). Tregs and CFSE- (1.5μM, Thermo Fisher scientific) labeled Tresp cells were mixed in a 1:2 (Treg: Tresp) ratio and stimulated with α-CD3/CD28 coated beads in a 1:2 ratio (beads: cells). CFSE dilution in Tresp was analyzed after 4 days of co-culture by flow cytometry.

### Immunofluorescence staining

Sections of formalin-fixed and paraffin-embedded tissue were cut in series at 4 μm, mounted on Superfrost Plus object glasses (ThermoFischer Scientific). The tissue was washed sequentially in xylene, ethanol and phosphate-buffered saline (PBS). Heat-induced epitope retrieval was performed by boiling sections for 20 minutes in DAKO buffer (pH 6) and cooled to room temperature before staining. Paired immunofluorescence staining was performed by applying a polyclonal rabbit anti-human CD3 antibody (1/50, A0452, Agilent) and rat IgG2a anti-human Foxp3 (clone PCH 101, 1/20, eBioscience) for 1 h at 37 °C, then raised in PBS. The secondary antibodies, Alexa Fluor 488 labeled donkey anti-Rabbit IgG (2 μg/ml, Molecular Probes, Eugene, OR USA) and a Cy3 conjugated AffiniPure donkey anti-rat IgG (H+L) (1/100, Jackson ImmunoResearch) were incubated for 1.5 h in room temperature and then washed in PBS. Sections where then incubated for 5 min at room temperature with Hoechst 33342 nucleic acid stain, and stained sections were mounted with ProLong Glass Antifade mountant (Molecular probes). Laser scanning confocal microscopy was performed by acquiring tile scans on an Andor Dragonfly equipped with a fusion stitcher. The Andor Dragonfly was built on a Nikon TiE inverted microscope, equipped with a 20x/0.75 NA air objective, and is equipped with a Zyla4.2 sCMOS camera.

## RESULTS

### Human SI Tregs are scarce but have strong suppressive activity

To investigate the Treg population in human SI, we obtained single cells from enzyme-digested mucosal tissue of histologically normal SI as well as from large intestine and peripheral blood mononuclear cells (PBMCs) for comparison. As reported earlier, we found that about 9% of memory CD4 T cells expressed Foxp3 in peripheral blood and in the lamina propria (LP) of colonic mucosa (Fig. 1a, b). However, only a median of 2% of LP CD4 T cells in the SI expressed Foxp3 (Fig. 1a, b). Immunostaining of tissue sections from normal SI and colon with antibodies to CD3 and Foxp3 showed a similar difference in Foxp3+ T cells (**Fig. 1c, d**). We detected very few Foxp3+ T cells in the SI epithelium (**Fig. 1c** and **Supplementary Fig. S1**) and further analysis therefore focused on Tregs in LP only. Most SI Foxp3+ CD4 T cells expressed a memory Treg phenotype being CD45RA^−^CD127^−^CTLA4^+^ (**Fig. 1e**). The transcription factor Helios was expressed by approximately 60% of Tregs in the SI, which was similar to that found in colon and PBMCs (**Fig. 1f, g**). Finding reduced fractions of Tregs in SI compared to blood and colon we next analyzed whether Treg numbers depended on the age of our different patient sample cohorts. When examining adults age 20 to 80 yrs (n=33), including both pancreatic cancer patients and donors for pancreas transplantation, we found a very slight but non-significant increase of both Helios+ and Helios-Tregs in the younger age groups (**Supplementary Fig. S2**).

**Fig. 1:**
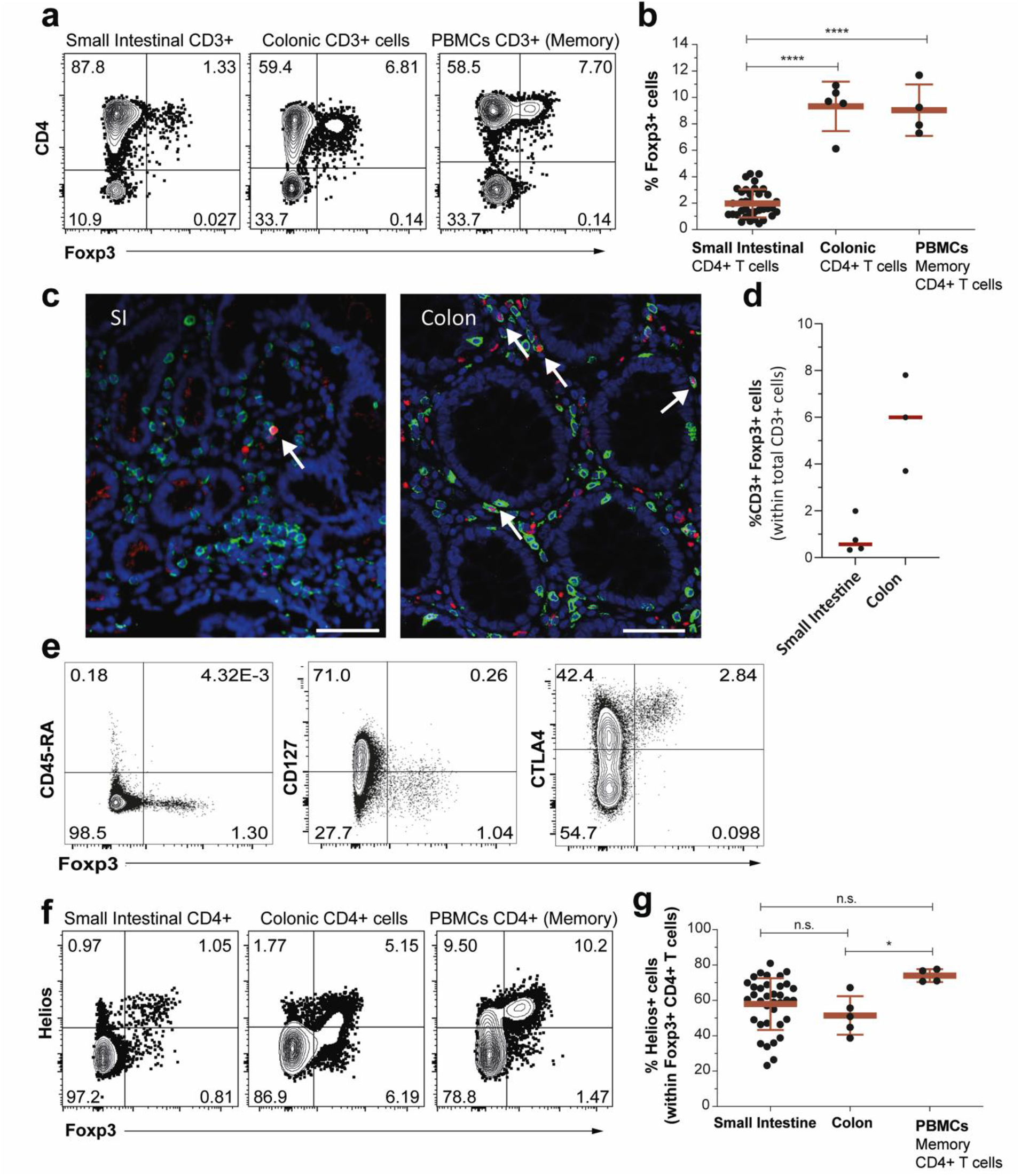
CD4 Tregs are scarce in the SI. **a)** Expression of CD4 and Foxp3 in CD3+ T cells from human SI, colon, and PBMCs (gated on CD45RO). Representative flow-cytometry dot plots (left) and compiled data (right) are shown showing the ) b) Percentage of Foxp3+ T cells in immunostained FFPE tissue sections from histologically normal duodenal and colonic mucosa. Representative confocal images stained for CD3 (green), Foxp3 (red), and hoechst (blue, nuclear stain) (left) and compiled data (right) are shown. Scale bars, 100 μm. **c)** Representative dot plots showing the expression of Foxp3, CD45RA, CD127 and CTLA-4 in SI CD4+ T cells. **d)** Expression of the transcription factors Helios and Foxp3 on total CD4+ T cells from SI, colon and CD45RO+ CD4+ T cells from PBMCs. Representative dot plots (left) and summary graphs (right) are shown. Data are given as mean ± SD and each dot represents one donor. ∗p < 0.05, ∗∗p < 0.01, ∗∗∗p < 0.001, ∗∗∗∗p < 0.0001 and non-significant (n.s.). One-way with Tukey’s multiple comparisons test.

To examine their suppressive activity, SI-derived Tregs were co-cultured with autologous naïve CD4 T cells from peripheral blood. High expression of CD25 is a hallmark of Tregs, however, surface CD25 was completely cleaved after enzymatic treatment during the sample preparation. In order to sort Tregs based on CD25 expression, we therefore cultured tissue-derived single cells for Tregs to regain CD25 expression. After overnight culture we found that 2-3% of CD4 T cells expressed CD25 and all of these were CD127^neg^. Further phenotyping showed that the vast majority CD25^+^CD127^neg^ expressed Foxp3 and more than half expressed Helios (**Fig. 2a**); a phenotype reminiscent of freshly isolated SI Tregs (**Fig. 1**). Sorted CD25^+^CD127^neg^ CD4+ Tregs were then stimulated with anti-CD3/CD28-conjugated beads for 48 h ^15^. Naïve CD4 T cells proliferated extensively in response to CD3/CD28 TCR stimulation, but when co-cultured with bead-stimulated SI-derived Tregs the proliferative response was strongly suppressed (**Fig. 2a**).

**Fig. 2:**
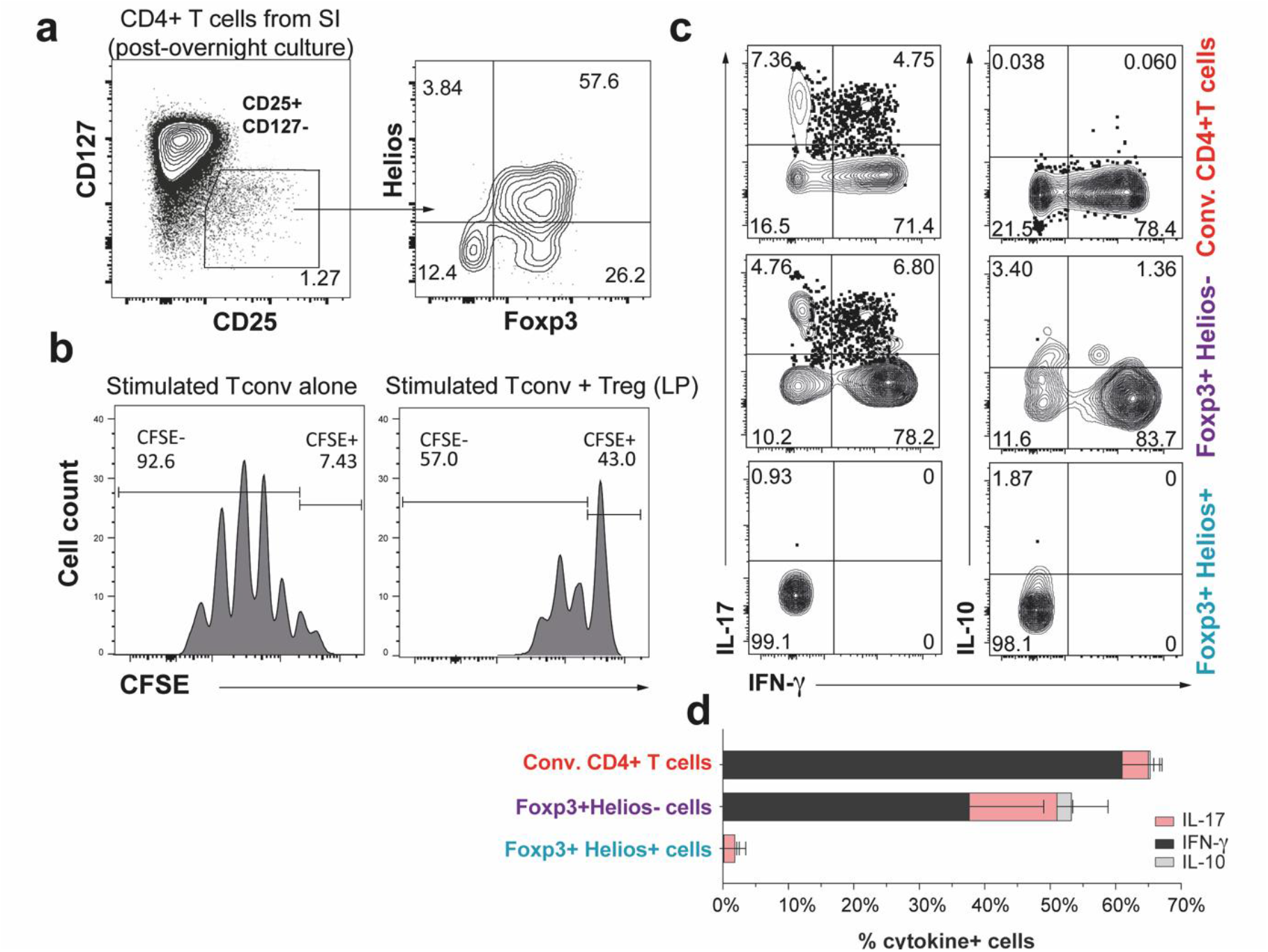
SI Tregs suppress autologous naïve T cells and produce cytokines. **a)** Dot plot showing the expression of CD127 and CD25 on SI CD4 T cells cultured overnight (left) and expression of Foxp3 and Helios gated in CD127^−^CD25^+^ CD4+ T cells (right). **b)** Histograms showing the expression of CFSE on blood-derived CD25^−^CD127^+^CD45RA^+^ CD4 T cells (CSFE-stained, named “responder cells”) stimulated with anti-CD3/CD28 beads for 4 days without (left) or with autologous stimulated SI-derived Tregs (CD25^+^CD127^−^CD4^+^ T cells). Representative of three independent experiments. **c)** Representative flow-cytometry dot plots and compiled data **(d)** of cytokine production after PMA and ionomycin activation of Foxp3-Helios− conventional, Foxp3+Helios+, and Foxp3+Helios− CD4 T cells in healthy SI.

We and others have shown that Helios+ and Helios− Tregs can be functionally distinguished by their ability to produce cytokines^16–18^. When SI-derived single cell suspensions were stimulated with PMA/ionomycin we found that Helios− Tregs, similar to conventional CD4 T cells, produced high levels of the proinflammatory cytokines IFN-γ and IL-17, and low levels of IL-10 (**Fig. 2b**). In contrast, these cytokines were almost undetectable in Helios+ Tregs (**Fig. 2b**).

Together, these findings show that Tregs are scarce in SI, but have strong suppressive capacity and displayed similar functional phenotypes as shown in blood and other mucosal tissues ^16-18^.

### Helios+ and Helios− Tregs display different replacement kinetics in transplanted SI

Studies in mice have shown that a large fraction of SI CD4 Tregs are dynamic cells with half-life of 4-6 weeks^7^. To directly assess the turnover rate of human SI CD4 Tregs we examined their replacement kinetics in transplanted duodenum. We obtained duodenal biopsies from type I diabetes patients undergoing pancreas transplantation with a duodenal segment (**Fig. 3a** and ^14^). Protocol biopsies were obtained from the duodenal graft and from the recipient (native) duodenum at 3, 6 and 52 weeks after transplantation. All recipient and donor pairs expressed different HLA class I alleles, allowing us to distinguish donor and recipient cells by flow cytometry (**Fig. 3b**) ^19–21^. This approach provides a unique possibility to study the lifespan of tissue-resident cells directly because there is no recruitment of donor cells from the circulation ^19, 21, 22^. Only patients without clinical and histological signs of rejection were included. We have previously reported that donor CD8 and CD4 T cells are very persistent in the transplanted tissue ^19, 20^. Here we show that the persistency of Helios− CD4 Tregs was similar to conventional Foxp3-CD4 T cells (**Fig. 3c, d**), whereas the lifespan of Helios+ CD4 Tregs was dramatically shorter with virtually no remaining donor-derived cells 1 yr post-transplantation (**Fig. 3c, d**). Interestingly, incoming (recipient) Tregs, constituted >8% of all recipient CD4 T cells at all three time points (**Fig. 3d**), whereas in the native (non-transplanted) duodenum both Treg subsets were stably present at steady-state levels (**Fig. 3d**).

**Fig. 3:**
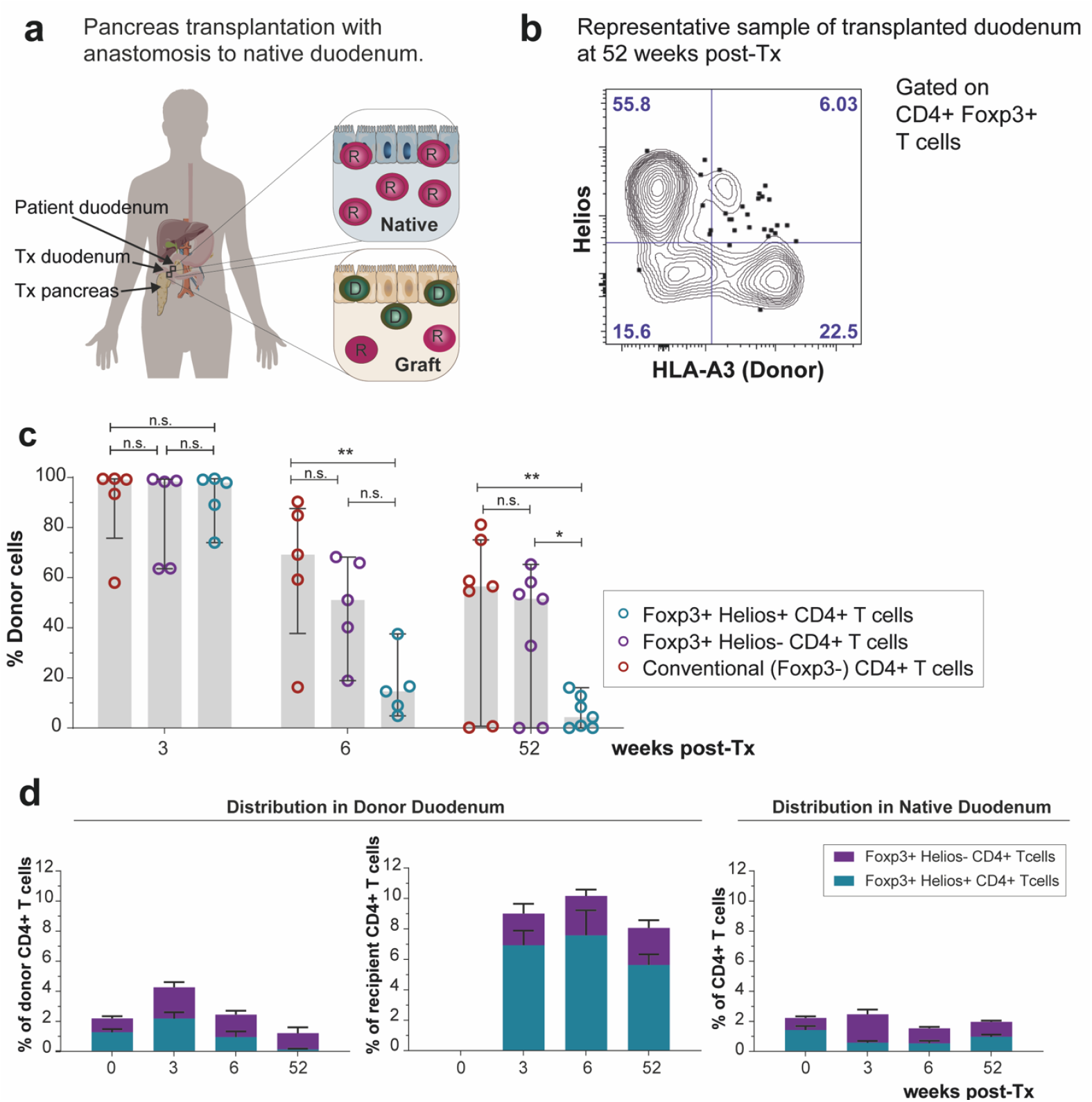
Helios+ Tregs are short-lived in transplanted small intestine. **a)** Depiction of pancreaticoduodenal transplantation procedure in type-I diabetic patients. **b**) Donor and recipient Foxp3+CD4 T cells in transplanted duodenum were distinguished based on disparate HLA class-I expression. Representative dot plot showing the expression of Helios and HLA-A3 on SI-derived Foxp3+ CD4 T cells 52 weeks after transplantation (donor HLA-A3^+^; recipient HLA-A3^−^). **c)** Percentage of donor cells in Helios+Foxp3+, Helios− Foxp3+, and conventional Foxp3-CD4 T cells in transplanted duodenum at 3, 6, and 52 weeks after transplantation as determined by HLA class I expression (as in b). Gray columns indicate median values and lines represent IQR. Each dot corresponds to an individual donor. Statistical analysis was performed using two-way ANOVA with repeated measures across subsets and Tukey’s multiple comparisons test of subset with time. n.s., not significant; *, p ≤ 0.05; **, P ≤ 0.01.**d)** Donor and recipient CD4 T cell subset distribution in donor (left) and recipient (native, right) and duodenum before (0) and after (3, 6 and 52) weeks post-transplantation. Mean with SEM is shown.

The difference in replacement kinetics between donor Helios− and Helios+ Tregs may have several explanations, including different egress rate into the draining lymphatics and/or different proliferative capacity. SI CD4 resident memory T (Trm) cells express receptors involved in tissue retention such as CD69,CD49a,and CD103, but not CCR7 that is involved in tissue egress ^20 23–25^. In order to compare the potential of the two Treg subsets to reside in or exit the tissue we analyzed the expression of these markers on CD4 T cells isolated from histologically normal SI. Interestingly, very few Helios-positive Tregs expressed CD69, CD49a and CD103 compared to CD4 T cells and Helios− Tregs, whereas the Helios-negative subset only differed from conventional T cells by less CD69+ cells (**Fig. 4a**). Both Treg subsets expressed more CCR7 than their conventional counterpart (**Fig. 4a**). To determine their proliferative capacity, we analyzed the expression of the proliferation marker Ki-67. While less than 2% conventional CD4 T cells expressed Ki67, approximately 10% of both of Treg subset expressed this marker (**Fig. 4b**).

**Fig. 4:**
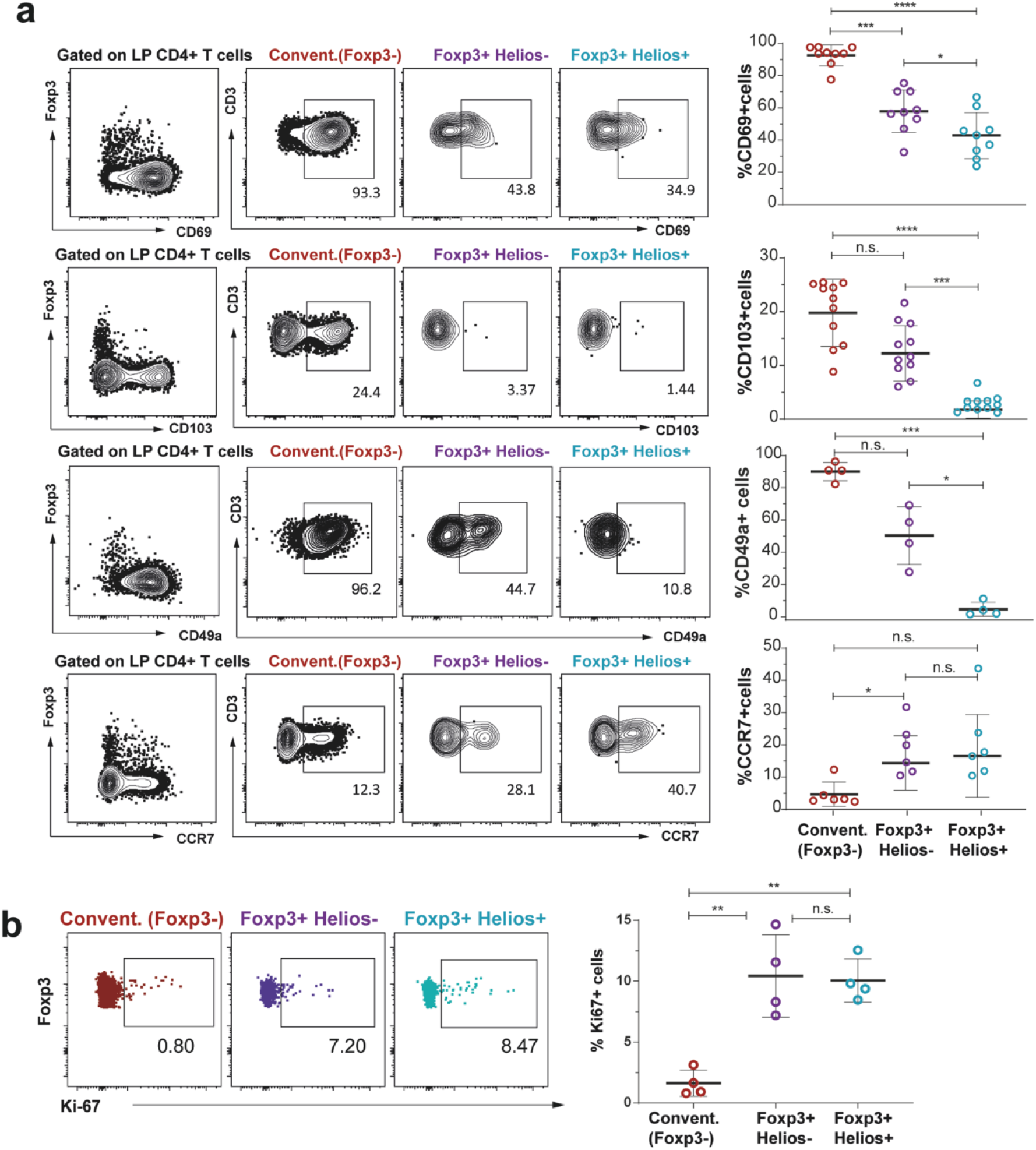
Intestinal Tregs present lower expression of Trm markers and more CCR7 than conventional CD4 T cells. **a) Expression of** CD69, CD103, CD49a, and CCR7 on SI-derived conventional CD4 T cells andHelios-and Helios+ Foxp3+ CD4 T cells. Representative dot plots (left) and compiled data (right) are shown. **b)** Expression of Ki67 on SI-derived CD4 and Foxp3+ CD4 T cells. Representative dot plots (left) and compiled data are shown (right). Data are shown as mean ± SD and each dot represents one donor. ∗p < 0.05, ∗∗p < 0.01, ∗∗∗p < 0.001, ∗∗∗∗p < 0.0001 and non-significant (n.s.). Repeated-measures one-way with Tukey’s multiple comparisons test.

Together, this demonstrates that Helios+ and Helios-Tregs have very different survival rates in the human SI. Helios-Tregs persist for more than one year whereas Helios+ Tregs are rapidly depleted from the transplanted tissue. This difference may to some extent be explained by their diverse expression of cell surface molecules involved in retention and egress of T cells.

### Both SI Treg subsets increase dramatically in active celiac disease

Next, we wanted to examine how SI Tregs respond in an inflammatory situation. To this end, we obtained SI biopsies from patients with newly diagnosed active celiac disease (CeD, n=14); a common chronic autoimmune inflammatory condition in human SI triggered by intake of dietary gluten in genetically predisposed individuals^26^. Analysis of tissue-derived single cell preparations from active CeD showed that nearly 20% of all CD4 T cells expressed Foxp3, of which approximately 60% co-expressed Helios (Fig. 5a, b). Immunostained tissue sections of the celiac lesion with antibodies to CD3 and Foxp3 showed similar high numbers (**Fig. 5c, d**). It has been shown that activated CD4 T cells may transiently upregulate Foxp3 ^27^. To determine whether Foxp3+ CD4 T cells in CeD are “bona fide” Tregs or activated T cells we performed three complementary experiments. First, we performed extensive phenotyping of CD4 T cells from the celiac lesion. **Fig. 6a** shows that both Helios+ and Helios-Foxp3+ CD4 T cells expressed a Treg phenotype, including high expression of CD95, CTLA-4, TIGIT, and CD39, and low expression of Trm-associated markers such as CD69, CD49a, CD103 and CD161. Conventional CD4 T cells, on the other hand, expressed high levels of Trm markers and low levels of Treg markers (**Fig 6a**). Secondly, we assessed the phenotype of CD3/CD28-stimulated CD4 T cells isolated from histologically normal SI. After stimulation there was 5 to 10-fold increased expression of Foxp3 and a reciprocal reduction of Helios expression, demonstrating that activated conventional CD4 T cells upregulated Foxp3 but not Helios (**Fig. 6b**). However, hierarchical clustering of immunophenotypic marker expression and principle component analysis (PCA) revealed that in vitro-activated Foxp3+ and Foxp3-CD4 T cells clustered together (**Fig 6c, d**), and were phenotypically very different from CeD-derived Foxp3+ CD4 T cells (**Fig. 6d**). Importantly, both CeD-derived Treg subsets clustered closely together with unstimulated Helios+Foxp3+ Tregs from normal SI. Together, these findings suggested that both Foxp3+ T-cell subsets in the celiac lesion were bona fide Tregs, and not activated CD4 T cells. Finally, we compared the capacity of CD4 T cells isolated from the celiac lesion and normal SI to produce cytokines after stimulating the cells PMA/ionomycin for 4 hours ^16–18^. Similar to T cells derived from normal SI (**Fig. 2 and 7**), CeD-derived Helios+ Tregs produced very low levels of IFNγ, IL-17 and IL-10 (**Fig. 7**). In contrast, Helios-Tregs produced significant levels of all three cytokines. Compared with Helios-Foxp3+ CD4 T cells from normal SI they produced significantly more IL-10, less IL-17, and similar amounts of IFNγ (**Fig. 7**). Conventional CD4 T cells from CeD produced slightly more IL-10 and similar levels of IL-17 and IFNγ compared to their counterparts from normal SI (**Fig 6e**). This 4h-stimulation did not alter the number of Foxp3+ T cells (**Supplementary Fig. S2**), ruling out the possibility that transient expression of Foxp3 by activated CD4 T cells were responsible for cytokine production observed by the Helios-Foxp3+ CD4 T cells.

**Fig. 5:**
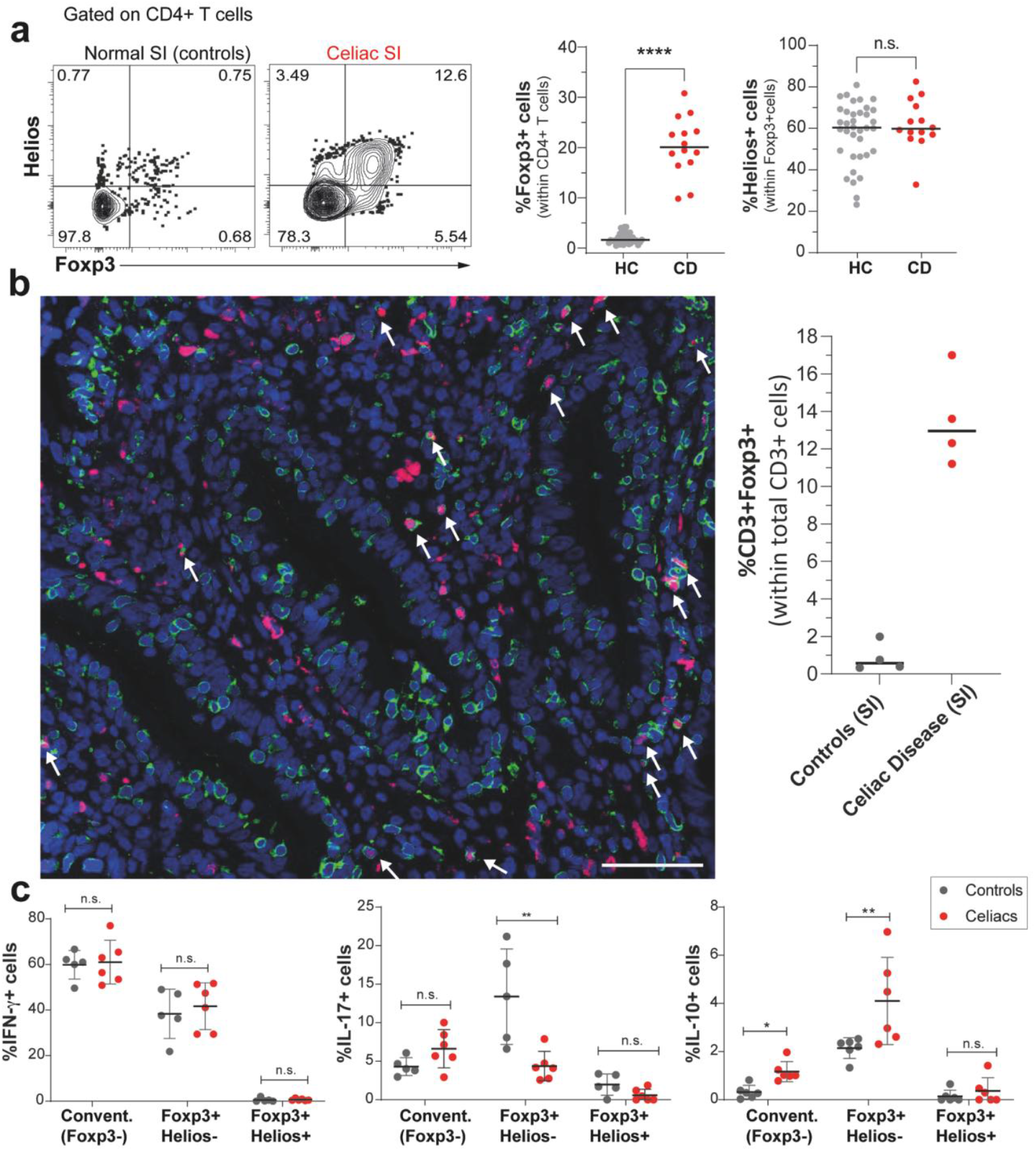
Helios+Foxp3+ and Helios-Foxp3+ CD4 T cells are dramatically increased in untreated celiac disease. **a)** Intranuclear expression of Foxp3 and Helios in CD4 T cells derived from normal SI and untreated celiac disease. Representative flow-cytometry dot plots (left) and summary graphs (right) are shown **b)** Percentage of Foxp3+ T cells in immunostained FFPE tissue sections from active celiac disease and controls. Representative confocal image from untreated celiac disease stained for CD3 (green), Foxp3 (red), and hoechst (blue, nuclear stain) (left) and compiled data (right). Scale bar, 100 μm **c)** Cytokine production in conventional Foxp3-Helios-, Foxp3+Helios-Foxp3+Helios+ CD4 T cells derived from normal and untreated celiac SI following 4h stimulation with PMA/ionomycin. Data are shown as mean ± SD and each dot represents one donor. ∗p < 0.05, ∗∗p < 0.01, ∗∗∗p < 0.001, ∗∗∗∗p < 0.0001 and non-significant (n.s.). Two-way ANOVA with Šídák's multiple comparisons test.

**Fig. 6:**
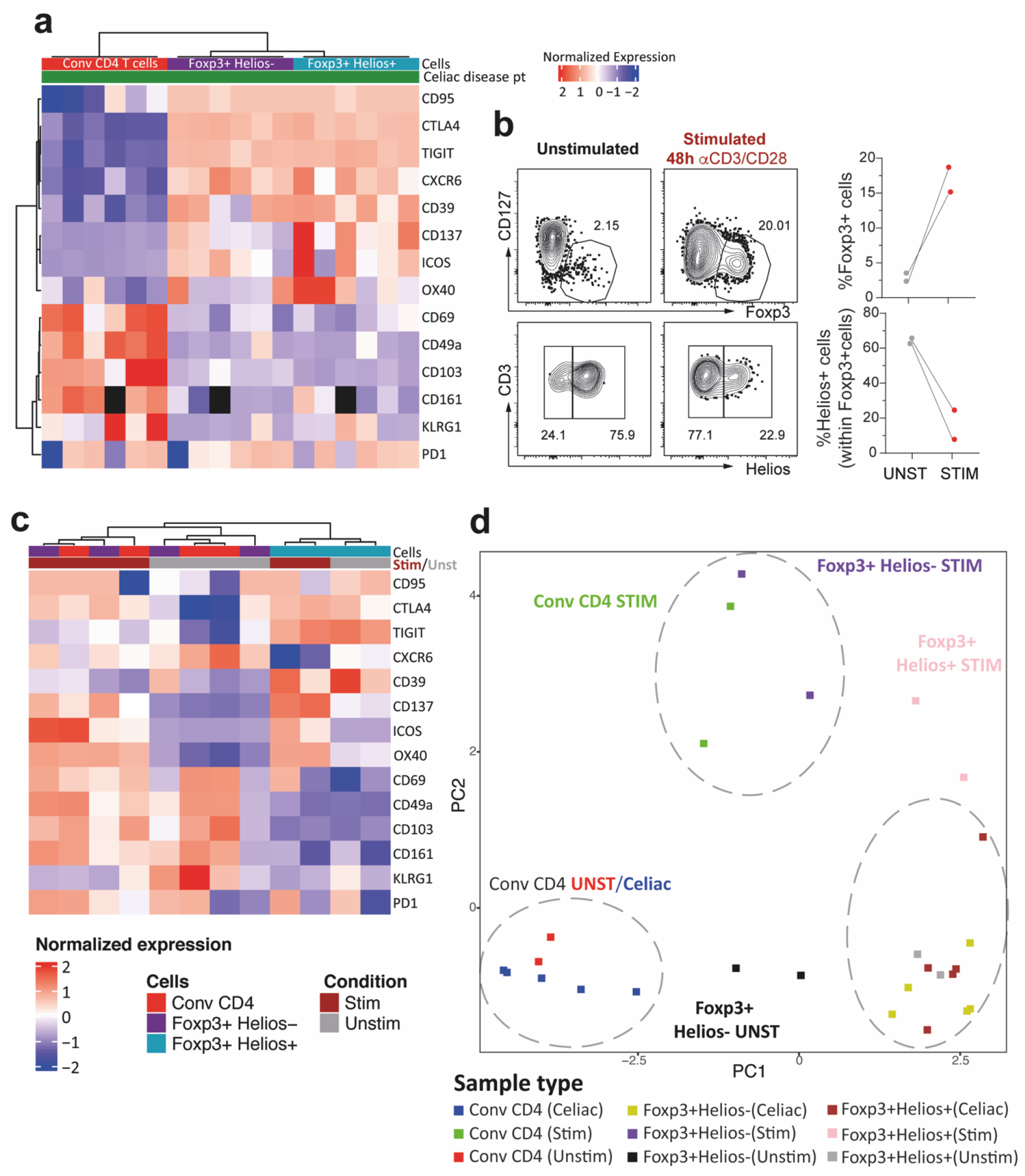
Foxp3+ CD4 T cells in untreated celiac disease are similar to steady-state “bona fide” Tregs, not conventional activated CD4+ T cells. **a)** Heatmap showing the expression intensity of various immunophenotypic markers on Foxp3+ Helios-, Foxp3+ Helios+ and conventional CD4 T cells isolated from biopsies of untreated celiac patients. Data are normalized by each row (for each single marker across all the patient samples) and hierarchically clustered. **b)** Intranuclear expression of Foxp3 and Helios in unstimulated and stimulated (48h with anti-CD3/CD28 beads) CD4 T cells from histologically normal SI. Representative flow-cytometry dot-plots (left) and compiled data (right) are shown‥ **c)** Heatmap showing the expression intensity of various immunophenotypic markers on unstimulated and stimulated Foxp3+ Helios-, Foxp3+ Helios+ and conventional CD4 T cells isolated from histologically normal SI. Data are normalized by each row (for each single marker across all the patient samples) and hierarchically clustered **d)** Principal Component Analysis (PCA) of Foxp3+ Helios-, Foxp3+ Helios+ and conventional CD4 T cells from untreated celiac patients and unstimulated/stimulated samples from histologically normal SI.

Taken together, these findings show that there was a large increase in both Helios+ and Helios-Tregs in the active celiac lesion. They were phenotypically and functionally very similar to their counterparts in normal SI mucosa, strongly suggesting that they represent bona fide Tregs and not activated T cells expressing Foxp3.

## DISCUSSION

Here we show that the human SI contains suppressive Foxp3+ CD4+ Tregs that comprise two subsets distinguished by their expression of the transcription factor Helios. Helios-Tregs show long-term tissue residency and produce substantial levels of both pro- and anti-inflammatory cytokines in response to stimulation, whereas Helios+ Tregs have a short half-life in the tissue and produce very low levels of cytokines. Both Treg subsets are scarce in the normal SI but are dramatically increased (approximately 10-fold) in lesions of active CeD.

Studies of Helios+ versus Helios- Tregs in humans have reported conflicting results. Helios-negative Tregs have been shown to be both more suppressive ^18^ and less suppressive ^28^ than Helios+ Tregs. Recently Lam et al. reported that Helios-KO and unedited Tregs had equivalent suppressive function, and that Helios is a marker, not driver, of Treg stability ^29^. We were not able to separate Helios+ from Helios- Tregs in our suppression experiments, but phenotypically, both subsets displayed a typical Treg phenotype, with high expression of markers such CTLA-4, TIGIT, CD95, and CD39.

It has been argued that activated T cells transiently upregulate Foxp3 under inflammatory conditions^27^, suggesting that a fraction of Foxp3+ cells in CeD-lesions may represent stimulated T cells. We found that conventional CD4 T cells isolated from normal SI upregulated Foxp3 when activated with anti-CD3/CD28. However, phenotypically these activated Foxp3+ T cells clustered together with activated conventional Foxp3-T cells, and far from both Helios- and Helios+ Foxp3+ T cells derived from lesions of active CeD, strongly suggesting all Foxp3+ CD4 T cells in active CeD were “bona fide” Tregs.

Importantly, SI Helios- and Helios+ Tregs showed very different replacement kinetics. Using unique human material of transplanted duodenum we demonstrated that Helios- Tregs were “long-lived” cells with a life span similar to Trm cells (>1 year), whereas Helios+ Tregs were replaced within weeks. Interestingly, the replacement kinetics of Helios+ Tregs is reminiscent of SI Tregs in mice that are induced by dietary antigens ^7^. Although the replacement kinetics was very different between the Helios+ and Helios- Treg subsets, their proportional numbers were similar in the steady-state and the CeD lesion – approximately 60% expressing Helios in both situations. Also, in the transplantation setting the relative proportions of Tregs among incoming CD4 T cells was remarkably stable through the 1-year period. Together, this suggested that, although the life span was very different, their tissue density appeared to be regulated by some common intrinsic and/or extrinsic mechanisms.

Helios− Tregs produced high levels of the proinflammatory cytokines IFN-γ and IL-17, whereas Helios+ Tregs did not. Several studies have shown that human Tregs with suppressive functions produce high levels of IL-17 ^30–32^, in particular in inflammatory diseases such as ulcerative colitis and Crohn’s disease ^30, 31^. Interestingly to this end, it has been reported that treatment with anti-IL-17 exacerbate inflammatory bowel disease ^33^, suggesting that IL-17 may have a protective function.

We found that SI Tregs constituted only 2% of all CD4 T cells in human adults, which is very low compared to 5-10% in other peripheral tissues (^16, 34^ and shown here). However, in an earlier study comparing tissue-resident Tregs in pediatric and adult organ donors it was found that the fractions of Tregs were age-dependent ^35^. In fact, the percentage of Tregs in the SI of infants under 2 yrs were 15-20% of all CD4 T cells, whereas the numbers in adult SI were similar to our findings. Based on studies in mice and earlier studies in infants, it is reasonable to believe that most SI Tregs are induced by food antigens ^7, 36^. This is compatible with high frequency of SI Tregs in infancy when a high number of new food items are introduced and furthermore suggest that maintenance of homeostasis (oral tolerance) to these antigens is less dependent on Tregs in adulthood. This seems to be different in the large intestine in which the fraction of Tregs is relatively high throughout life^37^. Several lines of evidence suggest that colon is more dependent of Tregs to avoid unwanted inflammation. Colitis is the most prevalent adverse effect when anti-CTLA-4 monotherapy is given ^12, 13^. Moreover, when IL-10 was selectively deleted in Foxp3+ Tregs in mice, the animals developed severe colitis and inflammation in the lungs, but no overt inflammation in the SI ^38^. It is well documented that the microbiota regulates the number and function of intestinal Tregs ^39^, and the high numbers of commensal bacteria in colon ^40^, may explain the high number of Tregs in this compartment of the intestine.

We found that the number of Tregs increased dramatically in active CeD and constituted almost 20% of total CD4 T cells. A role for Tregs in CeD has been disputed, but one study reported that the vast majority of gluten-specific CD4 T cells in blood of gluten-challenged CeD patients were Foxp3+ Tregs ^41^. However, these cells showed an impaired suppressive function. Other studies have shown a similar increase of Foxp3+ Tregs numbers in CeD, but found that the suppressive action of Tregs is reduced by local production of IL-15 ^42–44^. Increased numbers of Tregs are found in several inflammatory conditions in humans^45, 46^. However, their role in such conditions is still not clear. Interestingly, the relative number recipient (incoming) Tregs was much higher in the transplanted duodenum than the native duodenum at all time-points post-transplantation. We have previously reported that very few patients with pancreas-duodenal transplants show signs of rejection in the duodenal mucosa. It is tempting to speculate the high recruitment of Tregs plays a protective role in this transplantation setting.

Together, our results show that the human SI contains two Treg subsets with a very different life span and capacities to produce cytokines. Studies to explore their T-cell receptor repertoire and antigen specificities are awaited to further understand their functions in health and disease.

## Abbreviations used

IE: intraepithelial
LP: lamina propria
RPMI: Roswell Park Memorial Institute medium
SI: small intestine
Treg: regulatory T cell
Trm: resident memory T cell
Tx: pancreatic-duodenal transplantation (of diabetes mellitus patients)

## Acknowledgments

We are grateful to Kjersti Thorvaldsen Hagen, Hogne Røed Nilsen, and Kathrine Hagelsteen for excellent technical assistance; to the staff at the Endoscopy Units and the surgical staff; Christian Naper, Institute of Immunology, for providing HLA typing; Louise Fremgaard Risnes and Lene Støkken Høydahl, Institute of Immunology, for providing assistance with the celiac patients sample collection; the Confocal Microscopy and Flow Cytometry Core Facilities; all at Oslo University Hospital, Rikshospitalet.

## Author contributions

S.K. Chauhan, R. Bartolomé-Casado, E.S. Bækkevold, and F.L. Jahnsen conceived the project. S.K. Chauhan, R. Bartolomé-Casado and O.J.B. Landsverk, processed samples, designed and performed experiments, and analyzed data. S.K. Chauhan and R. Bartolomé-Casado prepared figures. S. Yaqub and R. Horneland coordinated recruitment of patients and collection of biopsies. S. Yaqub, R. Horneland, O. Øyen, and E.M. Aandahl performed surgery and provided samples. Knut E.A. Lundin performed endoscopy and provided endoscopic biopsies. S.K. Chauhan and F.L. Jahnsen wrote the manuscript. R. Bartolomé-Casado, O.J.B. Landsverk and E.S. Baekkevold contributed to writing the manuscript, E.S. Baekkevold, and F.L. Jahnsen supervised the study.

## References

1. Mowat AM. To respond or not to respond - a personal perspective of intestinal tolerance. Nat Rev Immunol 2018; 18:405–15.

2. Tanoue T, Atarashi K, Honda K. Development and maintenance of intestinal regulatory T cells. Nat Rev Immunol 2016; 16:295–309.

3. Sakaguchi S, Powrie F, Ransohoff RM. Re-establishing immunological self-tolerance in autoimmune disease. Nat Med 2012; 18:54–8.

4. Atarashi K, Tanoue T, Shima T, Imaoka A, Kuwahara T, Momose Y, et al. Induction of colonic regulatory T cells by indigenous Clostridium species. Science 2011; 331:337–41.

5. Campbell C, Dikiy S, Bhattarai SK, Chinen T, Matheis F, Calafiore M, et al. Extrathymically Generated Regulatory T Cells Establish a Niche for Intestinal Border-Dwelling Bacteria and Affect Physiologic Metabolite Balance. Immunity 2018; 48:1245–57 e9.

6. Lathrop SK, Bloom SM, Rao SM, Nutsch K, Lio CW, Santacruz N, et al. Peripheral education of the immune system by colonic commensal microbiota. Nature 2011; 478:250–4.

7. Kim KS, Hong SW, Han D, Yi J, Jung J, Yang BG, et al. Dietary antigens limit mucosal immunity by inducing regulatory T cells in the small intestine. Science 2016; 351:858–63.

8. Bacher P, Heinrich F, Stervbo U, Nienen M, Vahldieck M, Iwert C, et al. Regulatory T Cell Specificity Directs Tolerance versus Allergy against Aeroantigens in Humans. Cell 2016; 167:1067–78.e16.

9. Thornton AM, Korty PE, Tran DQ, Wohlfert EA, Murray PE, Belkaid Y, et al. Expression of Helios, an Ikaros transcription factor family member, differentiates thymic-derived from peripherally induced Foxp3+ T regulatory cells. Journal of immunology 2010; 184:3433–41.

10. Thornton AM, Shevach EM. Helios: still behind the clouds. Immunology 2019; 158:161–70.

11. Bennett CL, Christie J, Ramsdell F, Brunkow ME, Ferguson PJ, Whitesell L, et al. The immune dysregulation, polyendocrinopathy, enteropathy, X-linked syndrome (IPEX) is caused by mutations of FOXP3. Nat Genet 2001; 27:20–1.

12. Alissafi T, Hatzioannou A, Legaki AI, Varveri A, Verginis P. Balancing cancer immunotherapy and immune-related adverse events: The emerging role of regulatory T cells. J Autoimmun 2019; 104:102310.

13. Hodi FS, O’Day SJ, McDermott DF, Weber RW, Sosman JA, Haanen JB, et al. Improved survival with ipilimumab in patients with metastatic melanoma. N Engl J Med 2010; 363:711–23.

14. Horneland R, Paulsen V, Lindahl JP, Grzyb K, Eide TJ, Lundin K, et al. Pancreas transplantation with enteroanastomosis to native duodenum poses technical challenges--but offers improved endoscopic access for scheduled biopsies and therapeutic interventions. Am J Transplant 2015; 15:242–50.

15. Collison LW, Vignali DA. In vitro Treg suppression assays. Methods Mol Biol 2011; 707:21–37.

16. Ballke C, Gran E, Baekkevold ES, Jahnsen FL. Characterization of Regulatory T-Cell Markers in CD4+ T Cells of the Upper Airway Mucosa. PLoS One 2016; 11:e0148826.

17. Mercer F, Khaitan A, Kozhaya L, Aberg JA, Unutmaz D. Differentiation of IL-17-producing effector and regulatory human T cells from lineage-committed naive precursors. J Immunol 2014; 193:1047–54.

18. Raffin C, Pignon P, Celse C, Debien E, Valmori D, Ayyoub M. Human Memory Helios-FOXP3+ Regulatory T Cells (Tregs) Encompass Induced Tregs That Express Aiolos and Respond to IL-1beta by Downregulating Their Suppressor Functions. Journal of immunology 2013; 191:4619–27.

19. Bartolome-Casado R, Landsverk OJB, Chauhan SK, Richter L, Phung D, Greiff V, et al. Resident memory CD8 T cells persist for years in human small intestine. J Exp Med 2019; 216:2412–26.

20. Bartolome-Casado R, Landsverk OJB, Chauhan SK, Saetre F, Hagen KT, Yaqub S, et al. CD4(+) T cells persist for years in the human small intestine and display a TH1 cytokine profile. Mucosal Immunol 2021; 14:402–10.

21. Bujko A, Atlasy N, Landsverk OJB, Richter L, Yaqub S, Horneland R, et al. Transcriptional and functional profiling defines human small intestinal macrophage subsets. J Exp Med 2018; 215:441–58.

22. Landsverk OJ, Snir O, Casado RB, Richter L, Mold JE, Reu P, et al. Antibody-secreting plasma cells persist for decades in human intestine. J Exp Med 2017; 214:309–17.

23. Baekkevold ES, Yamanaka T, Palframan RT, Carlsen HS, Reinholt FP, von Andrian UH, et al. The ccr7 ligand elc (ccl19) is transcytosed in high endothelial venules and mediates T cell recruitment. J Exp Med 2001; 193:1105–12.

24. Cepek KL, Shaw SK, Parker CM, Russell GJ, Morrow JS, Rimm DL, et al. Adhesion between epithelial cells and T lymphocytes mediated by E-cadherin and the alpha E beta 7 integrin. Nature 1994; 372:190–3.

25. Shiow LR, Rosen DB, Brdickova N, Xu Y, An J, Lanier LL, et al. CD69 acts downstream of interferon-alpha/beta to inhibit S1P1 and lymphocyte egress from lymphoid organs. Nature 2006; 440:540–4.

26. Sollid LM. Coeliac disease: dissecting a complex inflammatory disorder. Nat Rev Immunol 2002; 2:647–55.

27. Wang J, Ioan-Facsinay A, van der Voort EI, Huizinga TW, Toes RE. Transient expression of FOXP3 in human activated nonregulatory CD4+ T cells. Eur J Immunol 2007; 37:129–38.

28. Bin Dhuban K, d’Hennezel E, Nashi E, Bar-Or A, Rieder S, Shevach EM, et al. Coexpression of TIGIT and FCRL3 identifies Helios+ human memory regulatory T cells. J Immunol 2015; 194:3687–96.

29. Lam AJ, Uday P, Gillies JK, Levings MK. Helios is a marker, not a driver, of human Treg stability. Eur J Immunol 2022; 52:75–84.

30. Hovhannisyan Z, Treatman J, Littman DR, Mayer L. Characterization of interleukin-17-producing regulatory T cells in inflamed intestinal mucosa from patients with inflammatory bowel diseases. Gastroenterology 2011; 140:957–65.

31. Jung MK, Kwak JE, Shin EC. IL-17A-Producing Foxp3(+) Regulatory T Cells and Human Diseases. Immune Netw 2017; 17:276–86.

32. Pesenacker AM, Broady R, Levings MK. Control of tissue-localized immune responses by human regulatory T cells. Eur J Immunol 2015; 45:333–43.

33. Pappu R, Rutz S, Ouyang W. Regulation of epithelial immunity by IL-17 family cytokines. Trends Immunol 2012; 33:343–9.

34. Sharabi A, Tsokos MG, Ding Y, Malek TR, Klatzmann D, Tsokos GC. Regulatory T cells in the treatment of disease. Nat Rev Drug Discov 2018; 17:823–44.

35. Thome JJ, Bickham KL, Ohmura Y, Kubota M, Matsuoka N, Gordon C, et al. Early-life compartmentalization of human T cell differentiation and regulatory function in mucosal and lymphoid tissues. Nat Med 2016; 22:72–7.

36. Karlsson MR, Rugtveit J, Brandtzaeg P. Allergen-responsive CD4+CD25+ regulatory T cells in children who have outgrown cow’s milk allergy. J Exp Med 2004; 199:1679–88.

37. Senda T, Dogra P, Granot T, Furuhashi K, Snyder ME, Carpenter DJ, et al. Microanatomical dissection of human intestinal T-cell immunity reveals site-specific changes in gut-associated lymphoid tissues over life. Mucosal Immunol 2019; 12:378–89.

38. Rubtsov YP, Rasmussen JP, Chi EY, Fontenot J, Castelli L, Ye X, et al. Regulatory T cell-derived interleukin-10 limits inflammation at environmental interfaces. Immunity 2008; 28:546–58.

39. Hegazy AN, Powrie F. MICROBIOME. Microbiota RORgulates intestinal suppressor T cells. Science 2015; 349:929–30.

40. Mowat AM, Agace WW. Regional specialization within the intestinal immune system. Nat Rev Immunol 2014; 14:667–85.

41. Cook L, Munier CML, Seddiki N, van Bockel D, Ontiveros N, Hardy MY, et al. Circulating gluten-specific FOXP3(+)CD39(+) regulatory T cells have impaired suppressive function in patients with celiac disease. J Allergy Clin Immunol 2017; 140:1592–603 e8.

42. Ben Ahmed M, Belhadj Hmida N, Moes N, Buyse S, Abdeladhim M, Louzir H, et al. IL-15 renders conventional lymphocytes resistant to suppressive functions of regulatory T cells through activation of the phosphatidylinositol 3-kinase pathway. J Immunol 2009; 182:6763–70.

43. Zanzi D, Stefanile R, Santagata S, Iaffaldano L, Iaquinto G, Giardullo N, et al. IL-15 interferes with suppressive activity of intestinal regulatory T cells expanded in Celiac disease. Am J Gastroenterol 2011; 106:1308–17.

44. Hmida NB, Ben Ahmed M, Moussa A, Rejeb MB, Said Y, Kourda N, et al. Impaired control of effector T cells by regulatory T cells: a clue to loss of oral tolerance and autoimmunity in celiac disease? Am J Gastroenterol 2012; 107:604–11.

45. Rajendeeran A, Tenbrock K. Regulatory T cell function in autoimmune disease. J Transl Autoimmun 2021; 4:100130.

46. van Herk EH, Te Velde AA. Treg subsets in inflammatory bowel disease and colorectal carcinoma: Characteristics, role, and therapeutic targets. J Gastroenterol Hepatol 2016; 31:1393–404.

