## Supplementary Material for "The human small intestine contains two subsets of regulatory Foxp3+ CD4+ T cells with very different life span and functional properties"

### Supplementary Table 1

| Target/Ab | Clone | Fluorophore(s) | Company | Reference |
|---|---|---|---|---|
| CD3 | OKT3 | BV510, PerCP-Cy5.5, Pe-Cy7, APC-eF780 | eBioscience | 47-0037-42 |
| CD4 | OKT4 | PerCP-Cy5.5, eF450 | Biolegend | 317428 |
| CD8 | SK1 | BV510, Alexa-Fluor488, PerCP Cy5.5, PE-Cy7 | Biolegend | 344716 |
| Foxp3 | 236A/E7 | PE-Cy7 | eBiosciences | 25-4777-42 |
| Helios | 22F6 | APC | eBiosciences | 17-9883-42 |
| CD25 | 2A3 | APC | BD Pharmingen | 340907 |
| CD127 (IL7-R) | Hil7r-m21 | BV605, BV711, PE | BD Horizon | 562662 |
| CTLA4 | L3D10 | PE | Biolegend | 349905 |
| CD45 | HI30 | BV510, Ax700, | Biolegend | 304036 |
| CD45 | 2D1 | APC-H7 | BD-Biosciences | 560178 |
| CD45-RA | HI100 | APC-eF780, PE-Cy7 | eBiosciences | 47-0458-42 |
| CD103 | Ber-ACT8 | BV605, PE-Cy7, FITC | Biolegend | 350218 |
| CD103 | Ber-ACT8 | BB515 | BD-Biosciences | 564578 |
| CD69 | FN50 | APC, BV605 | Biolegend | 310910 |
| CCR7 | G043H7 | PE | Biolegend | 353203 |
| HLA-A2 | BB 7.2 | PE | Abcam | ab79523 |
| HLA-A3 | GAP.A3 | FITC, APC | eBioscience | 11-5754-42 |
| HLA-B7 | BB7.1 | PE | Millipore | MAB1288 |
| HLA-B8 | REA145 | PE | Miltenyi | 130-118-960 |
| IFN-gamma | 4S.B3 | Ax488 | Biolegend | 502517 |

| | | | | |
|---|---|---|---|---|
| IL10 | JES3-9D7 | PE, BV421 | Biolegend | 564053 |
| IL17A | eBio64DEC17 | PE | eBiosciences | 12-7179-41 |
| Ki67 | B56 | Alexa 488 | BD-Biosciences | 558616 |
| CD49a | TS2/7 | PE | Biolegend | 328303 |
| KLRG1 | 13F12F2 | PE | eBiosciences | 12-9488-42 |
| CD28 | CD28.2 | PE | BD Horizon | 562976 |
| CD95 (FAS) | DX2 | PE | Biolegend | 305607 |
| CD39 | A1 | PE | Biolegend | 328207 |
| CTLA-4 | L3D10 | PE | Biolegend | 349905 |
| CD279 (PD-1) | EH12.2H7 | PE | Biolegend | 329905 |
| CD134 (OX40) | Ber-ACT35 (ACT35) | PE | Biolegend | 350003 |
| CXCR6 | K041E5 | PE | Biolegend | 356003 |
| CD161 | HP-3G10 | PE | Biolegend | 339903 |
| CD137 | 4B4-1 | PE | Biolegend | 309803 |
| TIGIT | MBSA43 | PE | eBiosciences | 12-9500-41 |
| CD278 (ICOS) | ISA-3 | PE | eBiosciences | 12-9948-41 |
| Isotype mouse IgG2a | MOPC-173 | PE | Biolegend | 400211 |
| Isotype mouse IgG1 | MOPC-21 | PE | Biolegend | 400414 |

**Supplementary Table 1.** Antibodies used in the study.

**SUPPLEMENTARY FIGURES:**

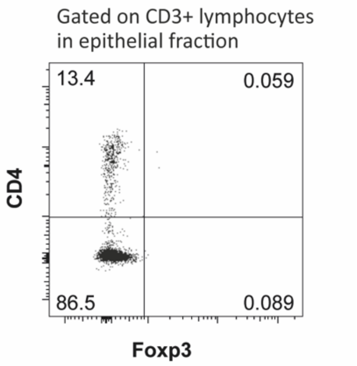


**Supplementary Fig. S1:** Representative dot plot showing the expression of Foxp3 and CD4 on CD3+ T cells in the epithelial fraction from normal small intestinal tissue.

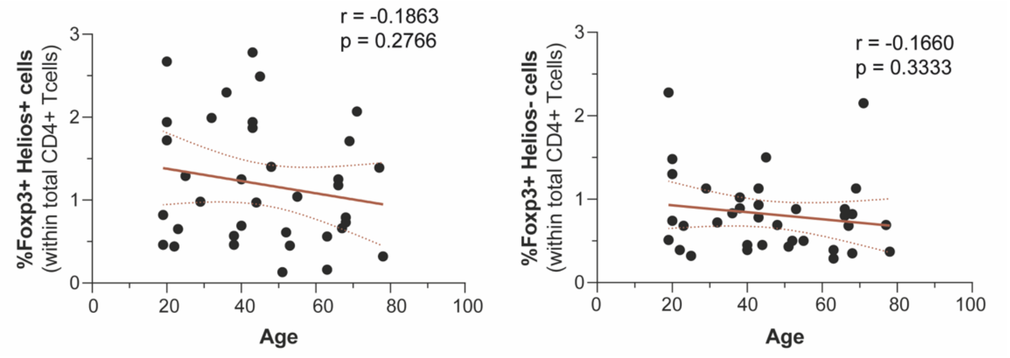


**Supplementary Fig. S2:** Distribution of Helios+Foxp3+ and Helios-Foxp3+ CD4 T cells in SI related to age.

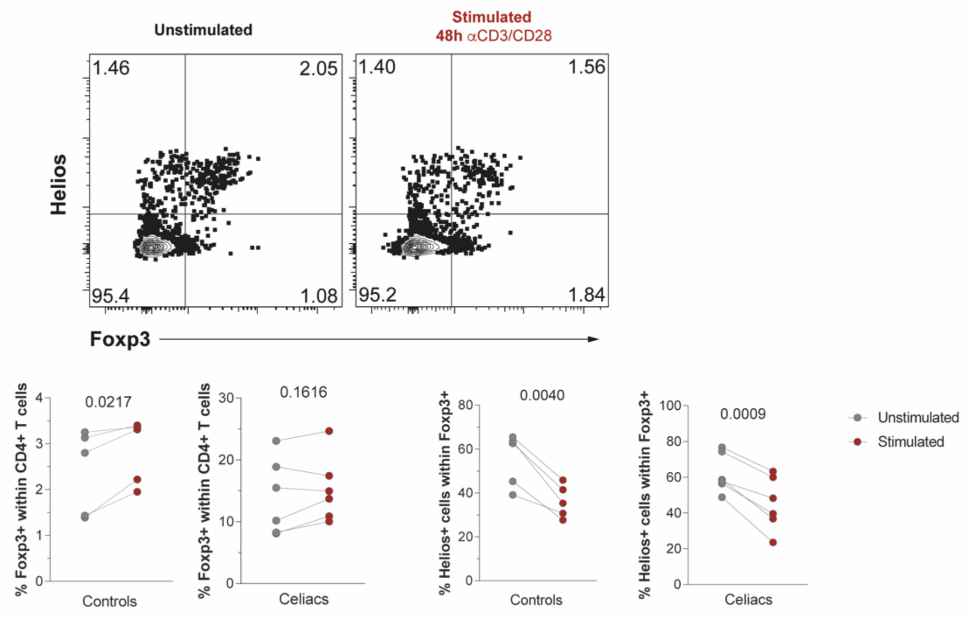


**Supplementary Fig. S3:** Expression of Helios and Foxp3 in unstimulated and stimulated CD4 T cells derived from active celiac disease lesions and normal control tissue.

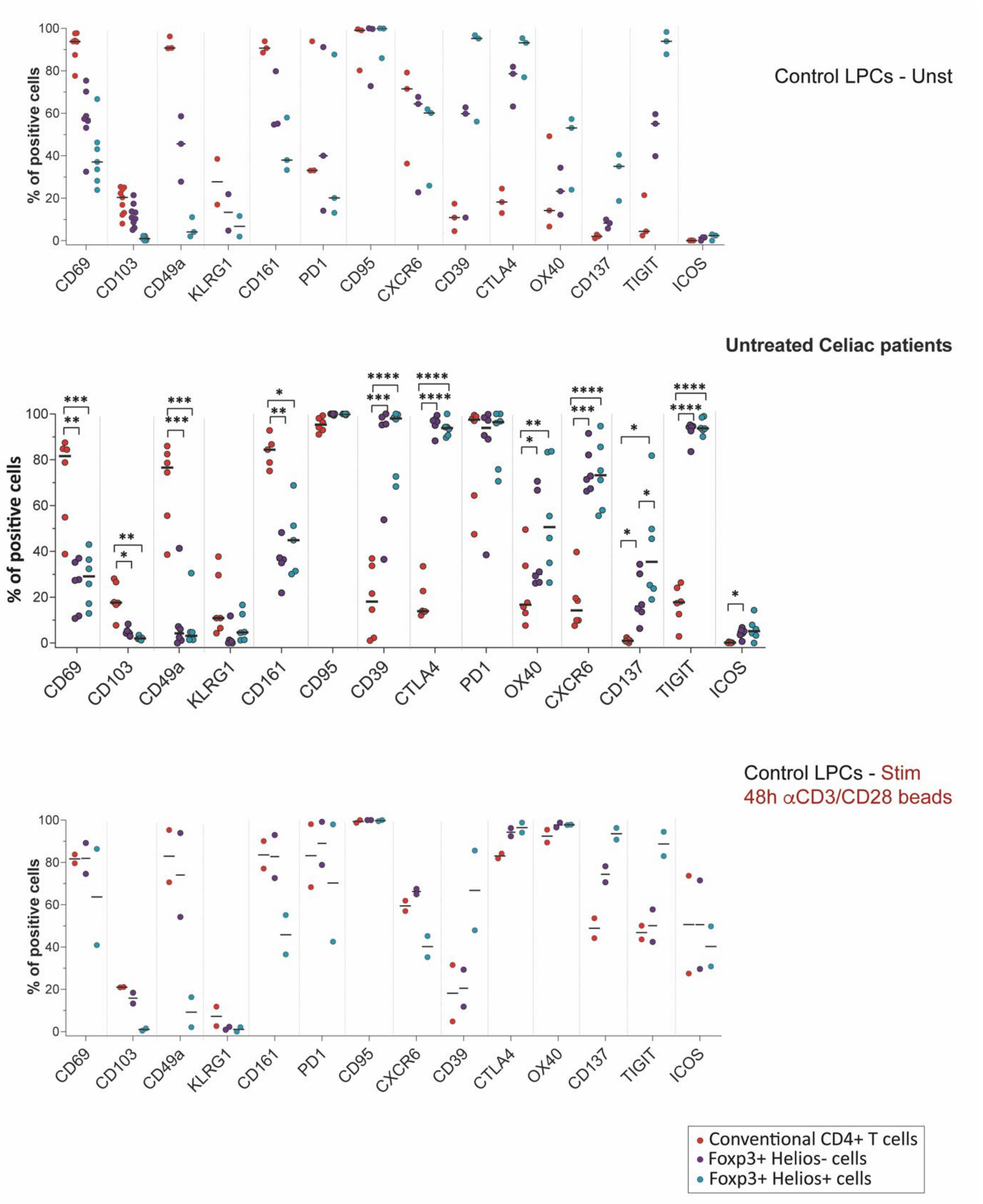


**Supplementary Fig. S4: Compiled expression data related to Figure 6**. Percentages of positive cells for the given markers on Foxp3+ Helios-, Foxp3+ Helios+ and conventional CD4 T cells isolated from biopsies of untreated celiac patients and unstimulated/48h stimulated control samples.
